# A high-throughput 3D kinetic killing assay

**DOI:** 10.1101/2023.04.11.536280

**Authors:** Renping Zhao, Archana K. Yanamandra, Bin Qu

**Affiliations:** Biophysics, Center for Integrative Physiology and Molecular Medicine (CIPMM), School of Medicine, Saarland University, Homburg, Germany; INM-Leibniz Institute for New Materials, Saarbrücken, Germany

**Keywords:** 3D matrix, kinetic, cytotoxicity assay, high throughput, high content imaging, translational immunology

## Abstract

In vivo, immune killer cells must infiltrate into tissues and search for their cognate target cells in 3D environments. To investigate the cytotoxic function of immune killer cells, there is currently a significant need for an in vitro kinetic assay that resembles 3D in vivo features. Our work presents a high-throughput kinetic killing assay in 3D that is a robust and powerful tool for evaluating the killing efficiency of immune killer cells, as well as the viability of tumor cells under in vivo-like conditions. This assay holds particular value for assessing primary human CTLs and NK cells and can also be applied to primary murine killer cells. By utilizing collagen concentrations to mimic healthy tissue, soft tumors, and stiff tumors, this assay enables the evaluation of cell function and behavior in physiologically and pathologically relevant scenarios, particularly in the context of solid tumors. Furthermore, this assay shows promise as a personalized strategy for selecting more effective drugs/treatments against tumors, using primary immune cells for individual patients to achieve improved clinical outcomes.

---

Immune killer cells, primarily natural killer (NK) cells and cytotoxic T lymphocytes (CTLs), play a crucial role in eliminating tumor cells. To carry out this function *in vivo*, these killer cells need to infiltrate into tissues and search for their cognate target cells in three-dimensional (3D) environments. Their killing efficiency is the result of a complex process that involves migration, recognition of target cells, formation of an immunological synapse, the release of lytic granules, and detachment from the target cells. To investigate the cytotoxic function of immune killer cells, various cytotoxicity assays have been developed, including assays that determine cytotoxicity at the endpoint [1-3] or in real-time [4]. However, these assays are typically performed in a two-dimensional (2D) setting. 3D *in vitro* assays, on the other hand, provide a physiologically relevant environment with defined features, making them powerful tools for investigating cell function and behavior [5-7]. In this work, we present a high-throughput kinetic killing assay in 3D, which allows for the quantitative analysis of the killing efficiency of CTLs or NK cells in real-time under physiologically and/or pathologically relevant conditions. This assay represents an important advancement in the study of immune cell function and could provide new insights into the mechanisms underlying immune-mediated tumor cell killing.

In this assay, the target cells are embedded in 3D collagen in a 96-or 384-well microplate, and settled at the well bottom in a 2D form after centrifugation to facilitate imaging. After solidification of collagen, killer cells are added from the top (Fig. 1A, step I). In this setting, the killer cells need to infiltrate into the 3D collage matrix (Fig. 1A, step II) and migrate to locate their target cells (Fig. 1A, step III). When the cognate target cells are identified, intimate contact is formed (Fig. 1A, step IV), inducing apoptosis in the target cells (Fig. 1A, step V). The target cell line (K562-pCasper) stably expresses an apoptosis reporter pCasper, which is a GFP-RFP FRET pair linked by a caspase cleavage site DEVD [8]. K562-pCasper target cells have been used to characterize the modes of NK cell killing in 3D collagen matrices [8]. When apoptosis is initiated, K562-pCasper cells lose their FRET signal, turning from orange-yellow to green, as shown in an exemplary experiment using NK cells in PBMCs as killer cells that directly recognize K562 cells (Fig. 1B, Movie 1, Fig. S1A). The killing dynamics were evaluated by determining the remaining living target cells, which were determined automatically by Imaris (Fig. 1C, Fig. S1B). This quantification can also be carried out semi-automatically by Fiji (TrackMate7) (Fig. S1C), and the results are comparable to those from Imaris (Fig. S1D). If a stable cell line expressing pCasper is not available, an alternative method is to stain target cells with CFSE and detect killing events in medium containing propidium iodide (PI) (Fig. 1D, Movie 2). This method distinguishes between destructed target cells (red) and living ones (green) (Fig. 1E, Fig. S2A). The quantification can be conducted using a similar pipeline established for pCasper (Fig. S2B-D). This CSFE/PI combination shows slower killing kinetics compared to pCasper-expressing cells (compare Fig.1C with Fig. 1F). This is likely because PI enters the cells after the integrity of the plasma membrane is compromised during apoptosis, which takes place after the initiation of apoptosis that can be directly detected by pCasper.

**Figure 1.**
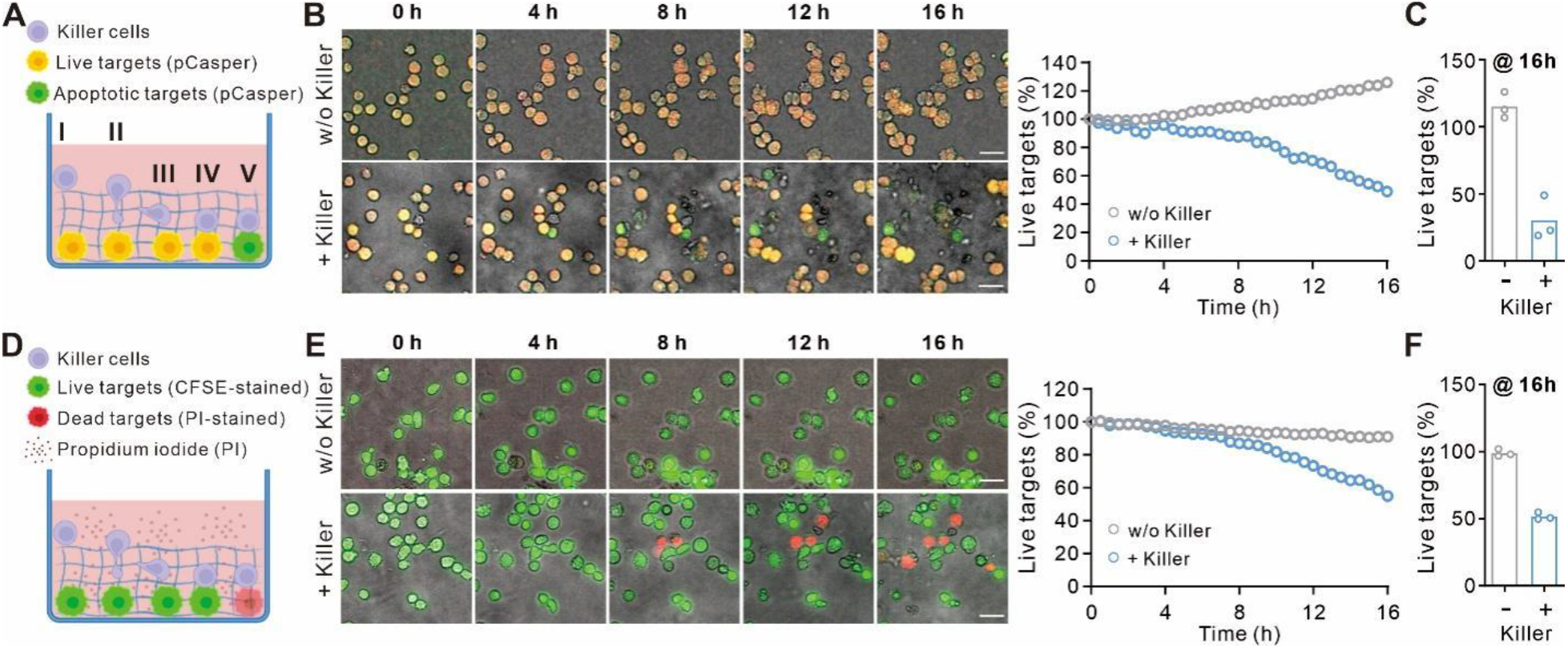
High content imaging based real-time 3D killing assay. (A) Schematic for killing-relevant events in this 3D killing assay. After adding from the top, killer cells first settle on the matrix (I), then infiltrate into the matrix (II) and migrate to look for their target cells (III). Upon recognition of target cells, immunological synapse is formed (IV). When apoptosis is initiated in target cells, they turn from orange-yellow color to green color (V). (B,C) K562-pCasper cells were embedded in bovine collagen I (2 mg/ml). PBMCs from healthy donors were used as killer cells. The killing events were visualized with a 10× objective every 30 min for 16 hours at 37°C with 5% CO_2_. (D-F) Using CFSE/PI to determine killing kinetics. K562 cells were stained with CFSE and then embedded in bovine collagen I (2 mg/ml). PI was present in the medium. PBMCs from healthy donors were used as killer cells. The killing events were visualized with a 20× objective every 30 min for 16 hours at 37°C with 5% CO_2_. Time lapse and the corresponding killing kinetics from one representative experiment out of three independent experiments is shown in B and E. Scale bars are 25 μm. The quantification at the end point (16 h) of all three experiments is shown in C and F. The effector (PBMCs) to target (E:T) ratio is 20:1. The schemes in A and D were created at biorender.com.

This assay allows for the testing of different concentrations of matrices. We compared the killing efficiency of CTLs in 2 mg/ml and 4 mg/ml collagen (Fig. 2A). The results show that the viability of target cells was not affected in either case (Fig. 2A, w/o CTLs), while the killing efficiency of CTLs was substantially reduced in denser collagen (4 mg/ml) compared to 2 mg/ml (Fig. 2A-B), which is consistent with our previous results [9]. In addition, this assay can be conducted in 384-well plates (Fig. S3) or using human collagen (Fig. S4).

**Figure 2.**
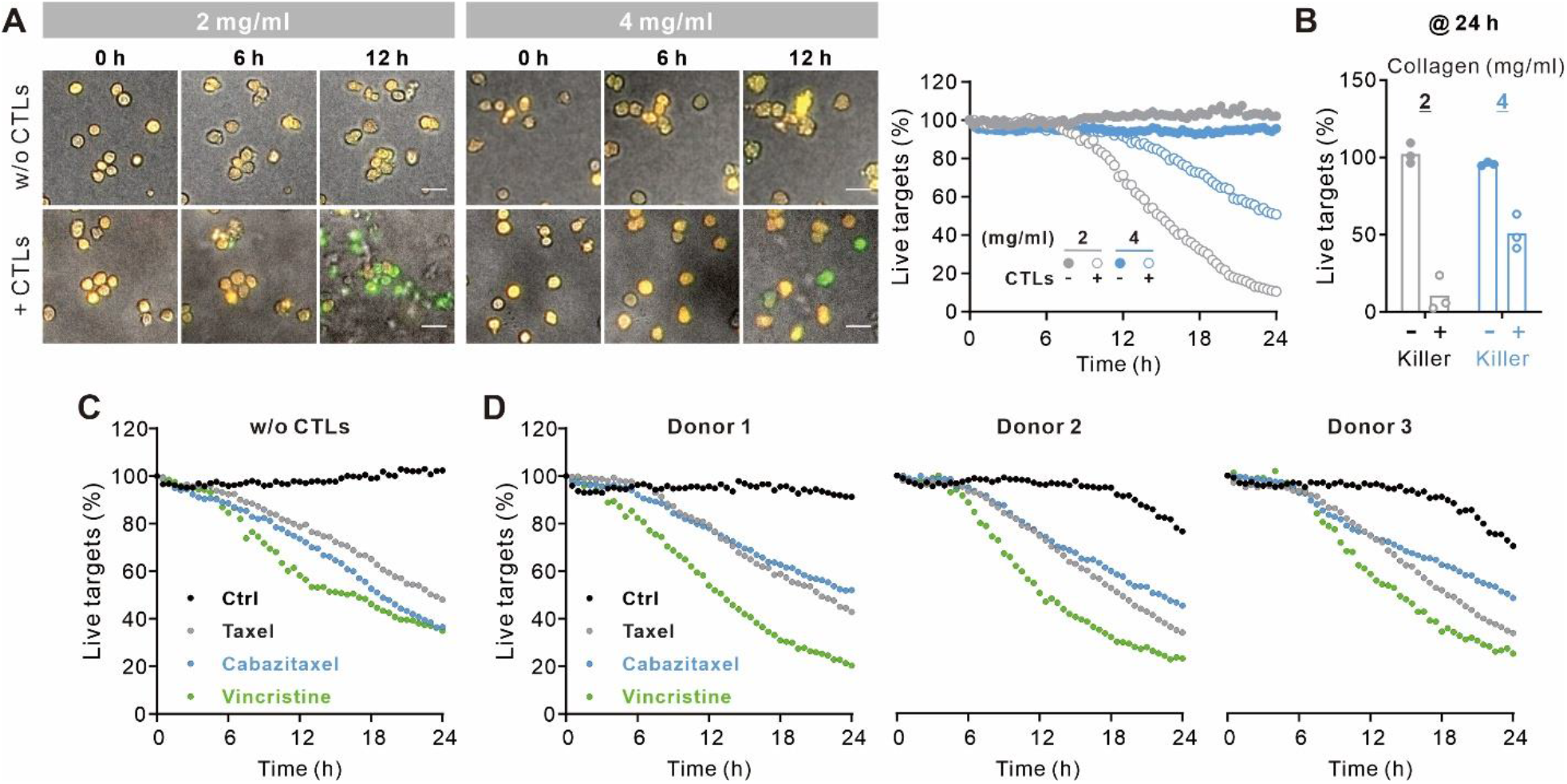
Applications of the real-time 3D killing assay. (A,B) Characterization of primary human CTLs killing efficiency in physiologically and pathologically relevant conditions. SEA/SEB pulsed NALM-6-pCasper cells were embedded in bovine collagen I (2 and 4 mg/ml mimicking healthy tissue and tumor [9], respectively. Bead-stimulated primary human CTLs were used as killer cells. Time lapse and the corresponding killing kinetics from one representative experiment out of three independent experiments is shown in A. Scale bars are 25 μm. The E:T ratio is 5:1. The quantification at the end point (24 h) of all three experiments is shown in B. (C,D) Drug screening with the real-time 3D killing assay. SEA/SEB pulsed NALM-6-pCasper cells were embedded in bovine collagen type I (2 mg/ml). Bead-stimulated primary human CTLs were used as killer cells. Taxel (0.4 μg/ml), Cabazitaxel (2 μg/ml), Vincristine (0.1 μg/ml) were present in the medium. PBS was used as control (Ctrl). The E:T ratio is 10:1.

This assay is not only useful for determining CTL/NK cell killing efficiency, but can also be used for high-throughput drug screening in 3D with or without killer cells. As an example, we tested three microtubule-targeting chemotherapeutic reagents, Taxel, Cabazitaxel and Vincristine. We treated SEA/SEB pulsed NALM-6-pCasper target cells embedded in collagen with these drugs at their clinically applied concentrations for 24 hours. We observed that these drugs induced apoptosis of tumor cells to comparable levels (Fig. 2C). When CTLs were added, the presence of these three drugs enhanced the overall killing of tumor cells relative to the control group (Fig. 2D). Interestingly, vincristine achieved better efficiency in eliminating tumor cells compared to Taxel or Cabazitaxel, particularly for Donor 1 and 2 (Fig. 2D). These findings suggest that these chemotherapeutic reagents can also influence the killing efficiency of immune killer cells, and that this method can be used to evaluate the possible clinical outcome of different drugs to optimize treatment strategies for patients.

In summary, this innovative 3D real-time killing assay presents a robust and powerful tool for assessing the killing efficiency of immune killer cells, as well as the viability of tumor cells under *in vivo*-like conditions. This assay holds particular value for evaluating primary human CTLs and NK cells and can also be applied to primary murine killer cells. By utilizing collagen concentrations to mimic healthy tissue, soft tumors, and stiff tumors [9], this assay enables the evaluation of cell function and behavior in physiologically and pathologically relevant scenarios, particularly in the context of solid tumors. Furthermore, this assay shows promise as a personalized strategy for selecting more effective drugs/treatments against tumors, using primary immune cells for individual patients to achieve improved clinical outcomes.

## Materials and methods please see Supporting Information

## Acknowledgments

We thank the Institute for Clinical Hemostaseology and Transfusion Medicine for providing donor blood; Carmen Hässig, Sandra Janku, Cora Hohxa, and Kethleen Seelert for excellent technical help; Markus Hoth for inspiring discussion and continuous support, as well as K562-pCasper cells (with Eva C. Schwarz); Xiangda Zhou and Shulagna Sharma for helpful discussion. This project was funded by the Deutsche Forschungsgemeinschaft (SFB 1027 A2 to B.Q.), Forschungsgroßgeräte (GZ: INST 256/429-1 FUGB) for ImageXpress, University of Saarland HOMFORexzellent grant (to R.Z.), Miniproposal of SFB1027 (to R.Z.).

## Ethical considerations

Research carried out for this study with human material (Leukocyte Reduction System Chambers from human blood donors) is authorized by the local ethic committee (Identification Nr. 84/15).

## Author contributions

R.Z. performed most experiments, conducted all the corresponding analyses, and created the figures; A.K.Y. conducted one experiment in Fig. 1C; B.Q. and R.Z. designed experiments; B.Q. conceptualized the study and wrote the manuscript with the assistance of R.Z.; all authors contributed to the writing of the manuscript and provided advice.

