## Supplementary Information for "A high-throughput 3D kinetic killing assay"

### Supporting Information

#### Materials and methods

##### Cell culture

Peripheral blood mononuclear cells (PBMCs) were isolated from healthy donors as described before [1]. Briefly, the content in Leukocyte Reduction System chambers was transferred with Hank's Balanced Salt Solution (Sigma-Aldrich) into 50 ml filtered tubes with Lymphocyte Separation Medium 1077 (Sigma-Aldrich) under the filter. After centrifugation (450 g for 30 mins at 4°C with the acceleration of 1 and deceleration of 0), the lymphocyte ring was collected. Then the remaining red blood cells were lysed with the erythrocyte lysis buffer (155 mM NH4Cl, 10 mM $KHCO_3$, 0.1 mM EDTA, pH=7.3). When PBMCs were used as killer cells, they were cultured in AIMV medium (ThermoFisher Scientific) with 10% Fetal Calf Serum (FCS) (ThermoFisher Scientific) and 50 ng/ml IL-2 (Miltenyi Biotec) for 2 days.

Human primary $CD8^+$ T cells were negatively isolated from PBMCs with Human $CD8^+$ T Cell Isolation Kit (Miltenyi Biotec) and then stimulated with Dynabeads™ Human T-Activator CD3/CD28 (ThermoFisher Scientific) for 2 days in AIMV medium supplemented with recombinant human IL-2 (17 ng/ml) and 10% FCS. Then the beads were removed from T cells and the stimulated T cells were cultured in AIMV medium supplemented with recombinant human IL-2 (17 ng/ml) and 10% FCS overnight prior to the experiments.

Human primary NK cells were negatively isolated from PBMCs using human NK Cell Isolation Kit (Miltenyi Biotec) and cultured for 2 days in AIMV medium supplemented with recombinant human IL-2 (50 ng/ml) and 10% FCS.

NALM-6-pCasper cells [2] and K562-pCasper cells [3] were established as described elsewhere. They were cultured in RPMI-1640 containing 10% FCS and 1% Penicillin-Streptomycin supplemented with puromycin (0.2 µg/ml) (VWR) for NALM-6-pCasper or G418 (1.25 mg/ml) (ThermoFisher Scientific) for K562-pCasper. All cells were cultured at 37°C with 5% $CO_2$.

##### SEA/SEB pulsing

NALM-6-pCasper cells ($1\times10^6$ cells/ml) were pulsed with staphylococcal enterotoxin A (SEA, 0.1 µg/ml) (Sigma-Aldrich) and SEB (0.1 µg/ml) (Sigma-Aldrich) at 37°C with 5% $CO_2$ for 40 mins in AIMV with 10% FCS. After centrifugation (200 g for 5 mins at room temperature), the pellet was resuspended in collagen for the 3D killing assay.

##### CFSE/PI staining

K562 cells ($1\times10^6$ cells/ml) were loaded with 5 µM of carboxyfluorescein succinimidyl ester (CFSE, ThermoFisher Scientific) in phosphate-buffered saline (PBS, ThermoFisher Scientific) at 37°C with 5% $CO_2$ for 20 mins. Afterwards, 5 times the original staining volume of RPMI-1640 containing 10% FCS was added and then the cells were incubated at 37°C with 5% $CO_2$ for another 5 mins. After centrifugation (200 g for 5 mins at room temperature), the pellet was resuspended in collagen for 3D killing assay. During the assay, PI (0.5 µg/ml, ThermoFisher Scientific) was present in the medium.

**High content imaging based real-time 3D killing assay**

Type I collagen (human or bovine as indicated, Advanced Biomatrix) was neutralized with NaOH as described elsewhere [4] and diluted to the required concentration with pre-chilled culture media. The cell pellet was resuspended in collagen and this cell/collagen mix was transferred in pre-chilled 96-well plates (Merck) with 18 µl for half-area and 40 µl for full-area 96-well plates. For 384-well plates (Merck), 1 mg/ml bovine collagen I was used and 10 µl of cell/collagen mixture for each well. the plate was centrifuged with 200 g for 5 mins at 4ºC and then flip the plate upside down and centrifuged again (200 g for 2.5 mins at 4ºC). The plate was incubated at 37 ˚C with 5% $CO_2$ for 1 hour to polymerize collagen. Afterwards, killer cells resuspended in AIMV/10% FCS were added from the top (200 µl for full-area 96-well plates, 100 µl for half-area 96-well plates, and 70 µl for 384 well plates) Concerning the number of target cells for each well, 25,000 for full-area 96-well plates, 12,500 for half-area 96-well plates, and 6,000 for 384-well plates. The following effector to target (E:T) ratios were used: 5:1 for NK:K562, 20:1 for PBMCs:K562, 5:1 or 10:1 for CTLs:NALM6.

The images were acquired by ImageXpress (Molecular Devices) with Spectra X LED illumination (Lumencor) at 37°C with 5% $CO_2$ every 20 mins for 12 to 24 hours. For K562-pCasper cells, LED 470/24 and Em 520/35 nm filter (Semrock) were used for the GFP channel, and LED 542/27 and Em 641/75 nm filter (Semrock) were used for the FRET channel. For CFSE/PI stained target cells, LED 470/24 and Em 520/35 nm filter (Semrock) were used to acquire signals from CFSE, and LED 542/27 and Em 641/75 nm filter (Semrock) were used to acquire signals from PI. A 20× S Fluor 0.75 numerical aperture (NA) objective (Nikon) or 10× S Fluor 0.5 NA objective (Nikon) was used as indicated in the figure legend.

**Quantification of killing kinetics with Imaris 9.6**

The images were first pre-processed with Fiji as follows: 1) the background of fluorescence was subtracted using 'Subtract Background'; 2) Only for CFSE-stained cells, the bleaching of CFSE was corrected by 'Bleach Correction' with Exponential Fit.

The TIFF files were converted to Imaris files (".ims") with ImarisFileConverter. The number of live target cells were determined using Imaris 9.6 as follows: 1) Open the file with Imaris; 2) Select 'Add new Spots'; 3) In the option 'Create' click 'Classify Spots' for Algorithm Settings and then click 'next'; 4) choose the GFP channel for pCasper cells or the CFSE channel for CFSE/PI-stained target cells, put an estimated XY diameter (80% of cell diameter, 10 µm for our cells and this diameter should be measured for each cell type) and tick 'Background Subtraction', then click 'next'; 5) choose 'Quality' for Filter Type and based on the first time point manually set the threshold as high as possible but still more than 95% of target cells can be detected with less than 1% false detection, then click 'next'; 6) select 'Fiter2D' for Filter Type, then choose 'Intensity Mean' for the corresponding channels (for pCasper cells, FRET for the x axis and GFP for the y axis); or choose 'Intensity Median' for the corresponding channels (for CFSE/PI-stained cells, CFSE for the x axis and PI for the y axis); 7) exclude the dead cells (for pCasper cells, gate the area under the slopingly positioned live cell population; for CFSE/PI-stained cells, gate the PI-positive population) then click ‘next’; 8) select ‘Statistics’ then click ‘Export All Statistics to File’ (the live target cells are labeled as ‘Class A’ in the exported file)..

**Quantification of killing kinetics with Fiji**

The images were first pre-processed with Fiji (background subtraction and bleach correction) as described above. The Plugin *TrackMate7* was used to determine the number of live target cells as follows: 1) open the file with Fiji, measure the diameter of the cells using “Straight line” tool;2) select ‘TrackMate’ from Plugins; 2) For LoG detector, detect in the GFP channel for pCasper cells or the CFSE channel for CFSE/PI-stained cells, put estimated object diameter (measured in the images), manually select Quality threshold as such that more than 95% of target cells can be detected with less than 1% false detection in Preview; 3) then click ‘next’ until the step of Display options, click ‘Spot’ to export the analysis.. Following criteria were used to identify live target cells: for pCasper cells FRET/GFP>0.4, for CFSE/PI-stained cells median fluorescence intensity of PI<23.

**Supplementary Figures**

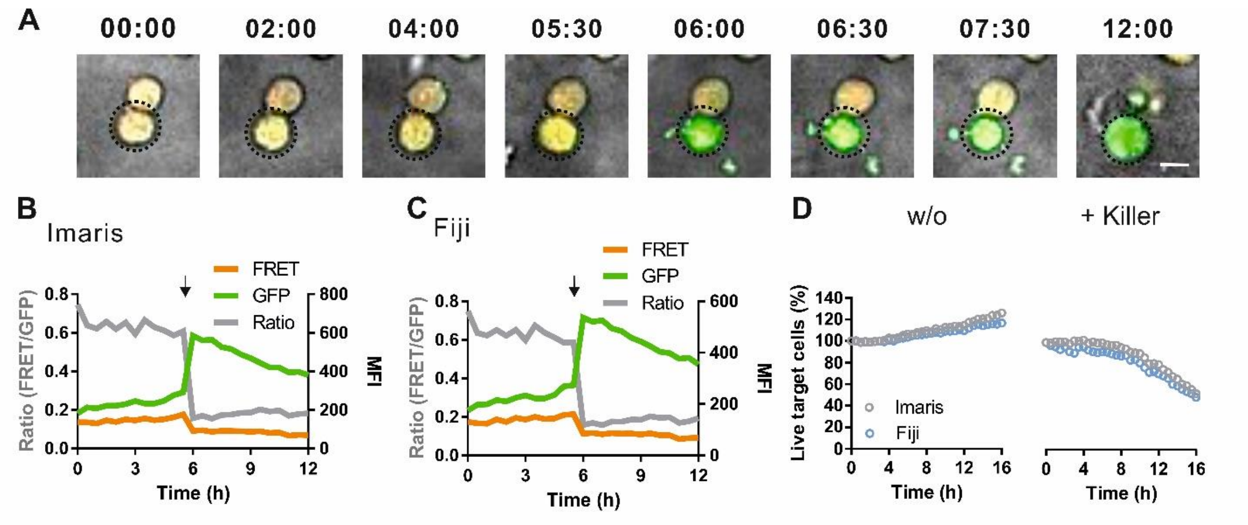


**Figure S1. Identification of apoptotic K562-pCasper cells.** K562-pCasper cells were embedded in bovine collagen I (2 mg/ml). PBMCs from healthy donors were used as killer cells. The killing events were visualized with a 10× objective every 30 mins for 16 hours at 37°C with 5% $CO_2$. Time lapse of an exemplary killing event is shown in **A**. GFP and FRET were shown in green and red, respectively. The target cell killed was highlighted with the black circle. The mean fluorescence intensity (MFI) of GFP signal and FRET signal as well as the FRET/GFP ratio are shown in the histograms analyzed by Imaris (**B**) or Fiji (**C**). The time point of cell death was pointed by black arrows. A comparison of quantification from Imaris and Fiji is shown in **D**. Scale bars are 10 µm.

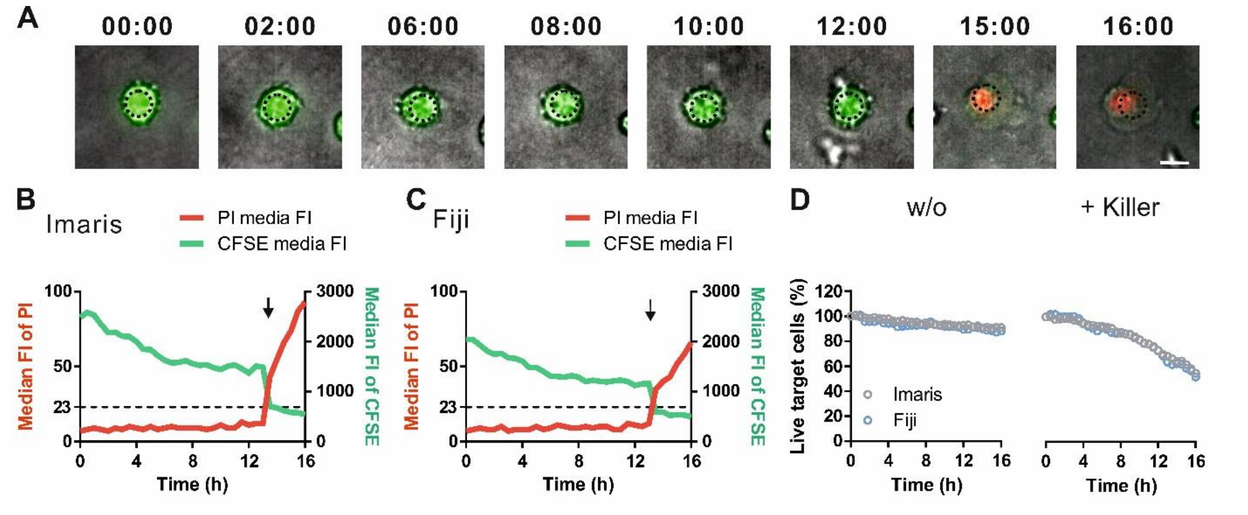


**Figure S2. Identification of apoptosis of CFSE-stained K562 cells along with PI.** CFSE-stained K562 cells were embedded in bovine collagen I (2 mg/ml). PBMCs from healthy donors were used as killer cells. The killing events were visualized every with a 20 × objective 30 mins for 16 hours at 37°C with 5% $CO_2$. Time lapse of an exemplary killing event is shown in **A**. CFSE and PI signals were shown in green and red, respectively. The target cell killed was highlighted with the black circle. The mean fluorescence intensity (MFI) of CFSE signal and PI signal as well as the CFSE/PI ratio are shown in the histograms analyzed by Imaris (**B**) or Fiji (**C**). The time point of cell death was pointed by black arrows. A comparison of quantification from Imaris and Fiji is shown in **D**. Scale bars are 10 µm.

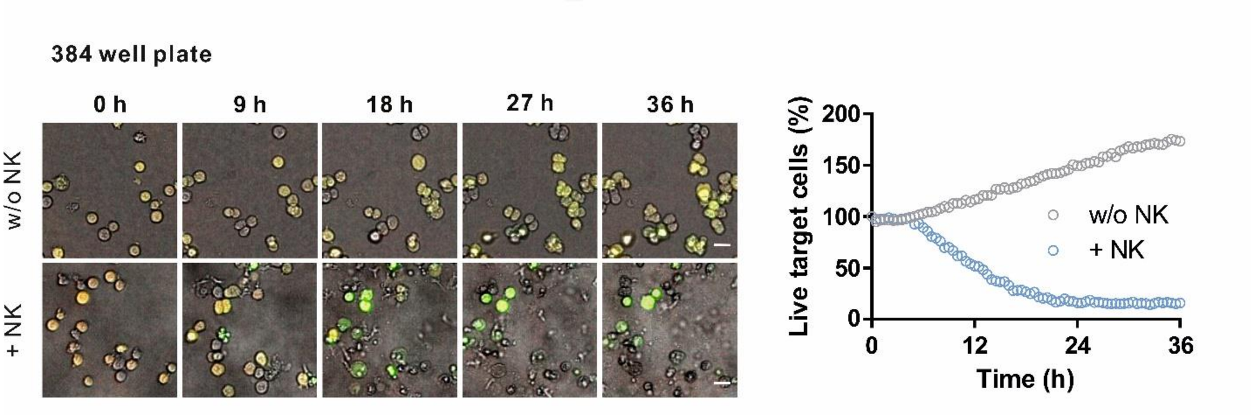


**Figure S3**. **High content imaging based real-time 3D killing assay conducted in a 384-well plate**. K562-pCasper cells were embedded in bovine collagen I (1 mg/ml). Primary human NK cells were used as killer cells. Images were acquired using ImageXpress with Spectra X LED illumination (Lumencor) with a 10× objective every 30 mins for 36 hours at 37°C with 5% $CO_2$. Scale bars are 20 µm. One representative example out of three independent experiments is shown.

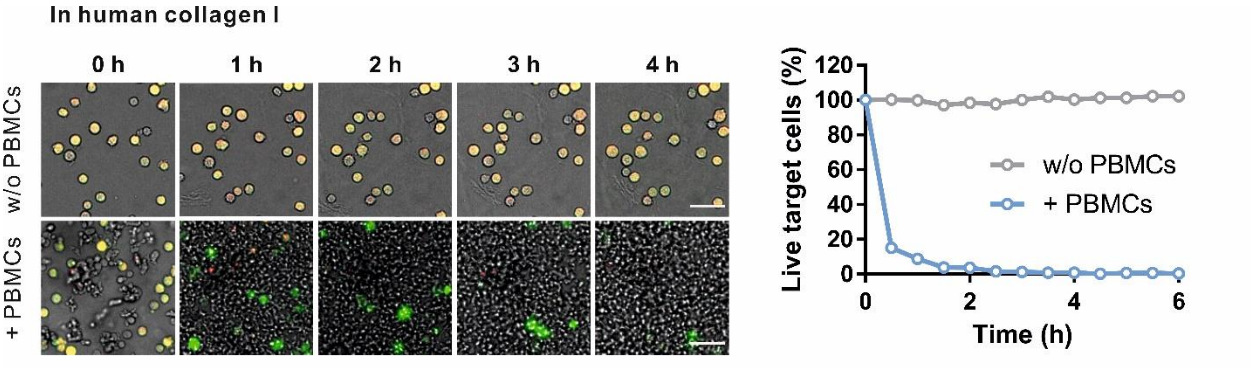


**Figure S4**. **High content imaging based real-time 3D killing assay with human collagen.** K562-pCasper cells were embedded in human collagen I (2 mg/ml). PBMCs were used as killer cells. Images were acquired using ImageXpress with Spectra X LED illumination (Lumencor) every 30 mins for 6 hours at 37°C with 5% $CO_2$. Since killing was completed by 6h, quantification of the first 6 hours is shown. Scale bars are 25 µm. One representative example out of three independent experiments is shown.

#### Movie legends

**Movie 1. Identification of apoptotic K562-pCasper.** K562-pCasper cells were embedded in bovine collagen I (2 mg/ml). PBMCs from healthy donors were used as killer cells. The entire field is shown, from which the region of interest was cropped and shown as time lapse in Fig. 1B. The original recording is shown on the left (green: GFP, red: FRET). The apoptotic cells identified with Imaris is shown on the right. Scale bars are 80 µm.

**Movie 2. Identification of CFSE-stained K562 cells in presence with PI.** CFSE-labeled K562 cells were embedded in bovine collagen I (2 mg/ml). PBMCs from healthy donors were used as killer cells. PI is present in the medium to define dead target cells. The entire field is shown, from which the region of interest was cropped and shown as time lapse in Fig. 1D. The original recording is shown on the left (green: CFSE, red: PI). The apoptotic cells identified with Imaris is shown on the right: Scale bars are 80 µm.
